# A systematic review of the effects of recreation on mammals and birds in mountains: Insights and future research directions

**DOI:** 10.1101/2022.09.08.507092

**Authors:** Adrian Hochreutener, Reto Rupf, Catherine Pickering, Claudio Signer

## Abstract

Mountainous areas are popular destinations for outdoor recreation, which can have environmental impacts on wildlife. We assessed research studies about the impacts of recreation on mammals and birds in montane, subalpine and alpine zones using a systematic literature review methodology to identify trends and gaps. We found that research on this topic so far has concentrated on specific regions, seasons, infrastructure, activities and taxa. Most of the 67 articles revived were from Europe (52%) or Northern America (37%) and mainly from subalpine habitats (49%), focused on recreation infrastructure (51%) and mainly done either in summer (47%) or winter (25%). Research was not taxonomically representative but focused on cervids (*Cervidae*, 21%), bovids (*Bovidae*, 17%) and grouse (*Phasianidae*, 8%). It included few species of high conservation value. Almost all research (91%) found significant effects, which were predominantly negative (82%). Infrastructure, such as trails, had the most evidence for negative effects, followed by activities such as hiking and backcountry skiing. Much of the research looked at impacts at individual (42%) or population level (40%) responses, such as changes in behaviour or reductions in habitat, with limited research on communities (7%) or for popular activities such as mountain biking. We invite researchers to make use of emerging technologies, such as remote sensing, and to address research gaps including more regions, taxa and activities. Utilizing current research, land managers can implement more evidence-based strategies to minimise impacts of recreation and mitigate human-wildlife conflicts.

## 1 Introduction

Outdoor recreation is popular in many mountainous areas (1). The activities comprise winter sports such as skiing, ice climbing or mountaineering, while in the snow-free seasons hiking, mountain biking and trail running are popular (2). The ecosystems in mountainous areas are often biodiversity hotspots and/or protected areas (3) that conserve a diversity of mammals and birds (4,5). The popularity of mountainous areas for recreation reflects broader social trend (6,7), such as more leisure time, (8) and more people wanting to living in or traveling to such areas (9). There, the impacts of recreation may be particularly acute as climatic conditions can be severe, with limited favourable periods for growth and reproduction (10). Also, in the past many of those areas experienced less human disturbance compared to lowlands (11), but this is shifting, including rapid changes in climate conditions (12). As a result there is growing pressures on the biodiversity of these habitats and the animals inhabiting them (13–16). Mammals and birds in particular have been focus of more research than other taxa (17) and many of them are endemic, threatened (18,19) and/or keystone species (23). Despite occurring in relatively remote areas, they have been adversely affected by a range of human activities and land uses in many places (5). But which species are affected - where, how, and by what types of recreation activities and infrastructure? Such information is crucial for those responsible for maintaining the conservation values and other ecosystem services of mountainous area while facilitating appropriate recreational use (1).

Research examining the ecological effects of recreation and tourism has documented a diversity of negative impacts on fauna, flora, soils and aquatic systems (20) as well as ways in which some impacts can be mitigated, minimised or even eliminated with appropriate management (21–23). Reflecting the importance of such research, reviews have assessed impacts of specific activities (1,24–26), for a range of ecosystems (30,34, 38,39) and types of biota (18-20) including in mountainous areas (18,29). For example, Sato et al. (18) in 2013 reviewed 35 years of research consisting of 31 articles that assessed the impacts of winter recreation and ski resorts on subalpine and alpine wildlife. They found changes in species richness, abundance and diversity associated with these types of tourism developments and activities. Larson et al. (30) in 2016 reviewed 274 articles on the impacts of recreation on wildlife, finding that non-consumptive recreation often had negative effects with snow-based activities more harmful than summer activities. Steven et al. (31) in 2011 summarized research on the impacts of nature-based recreation on birds using data from 69 articles, and found a wide range of negative impacts from different types of recreation. These three reviews all highlighted major gaps in recreation research including research from Africa, Asia and South America, for emerging recreation activities, for specific biota and research in spring and autumn. But how has the research community responded post these reviews?

To assess recent research, also about the gaps identified in previous reviews, we examined the impact of outdoor recreation on mammals and birds in mountainous areas. Using a systematic literature review methodology, we examined: 1) When, where and how has research been conducted? 2) Was research effort vs. demand (based on mountainous area per country) proportional? 3) Which activities have been assessed? 4) Which mammals and birds were studied and is research effort proportional to species diversity? 5) How did mammals and birds respond to specific activities and infrastructures? 6) What recommendations have been made on how to minimize or mitigate impacts? 7) Which research gaps remain?

## 2 Methods

### 2.1 Literature search strategy

We conducted a systematic quantitative literature review using the methods outlined in Pickering and Byrne (32) and Pickering et al. (33) including using the PRISMA protocol for systematic reviews (34) to create a database of peer-reviewed literature about impacts of outdoor recreation in mountainous areas on mammals and birds. Such reviews not only summarise current knowledge (32,33), they facilitate evidence based decisions for land managers (35) and can quantify gaps between the demand for research and its supply (36). By using a similar approach as previous key reviews (18,30,31) we could make direct comparisons, while extending the scope of those reviews to provide novel insights including if some of the gaps they identified are starting to be addressed. Specifically our review covered research from the beginning of 2005 until the end of 2019 on outdoor recreational activities and infrastructures (trails, resorts, ski lifts, recreational homes etc), in montane, subalpine and alpine zones, but excluding some types of consumptive activities such as hunting, fishing, berry- or mushroom picking (37).

The first stage of the systematic review involved searching the Web of Science core collection on 10^th^ May 2020 using a specific search string (Table 1) for the title, abstract, author and keywords of articles in English that covered relevant habitat, activities, responses and biota. The search string was developed iteratively during the pilot stage of the research and incorporated terms used in the titles, abstracts and keywords of key known literature and in the related reviews. We checked the effectiveness of the search terms by checking if the identified relevant articles from the previous reviews in overlapping search periods, as well as consulting experts and included additional relevant articles not identified by our search string (Fig 1: additional sources).

**Table 1:** Search string used to identify articles in Web of Science describing the effects of non-consumptive outdoor recreation on mammals and birds in mountainous areas.

| Topic | Search string |
| --- | --- |
| Habitat | (alpine OR subalpine OR mountain* OR montane OR highland*) |
| Recreation | AND ((recreation AND ecology)<br>OR “winter tourism” OR “snow tourism” OR “summer tourism”<br>OR “wildlife tourism” OR “nature tourism” OR “nature activit*”<br>OR “nature recreation” OR “winter recreation” OR “summer recreation”<br>OR “human recreation”<br>OR “outdoor activit*” OR trail* OR visitor* OR hik* OR walk* OR<br>“mountain bik*” OR mountaineering OR climb* OR skiing OR<br>(backcountry AND ski*)) |
| Response | AND (“habitat use” OR “human disturbance” OR disturbance OR stress<br>OR impact OR behaviour OR activity) |
| Biota | AND ((bird* OR avifauna) OR (mammal* OR ungulate*) OR wildlife)<br>NOT plant*. |

**Table 2:**
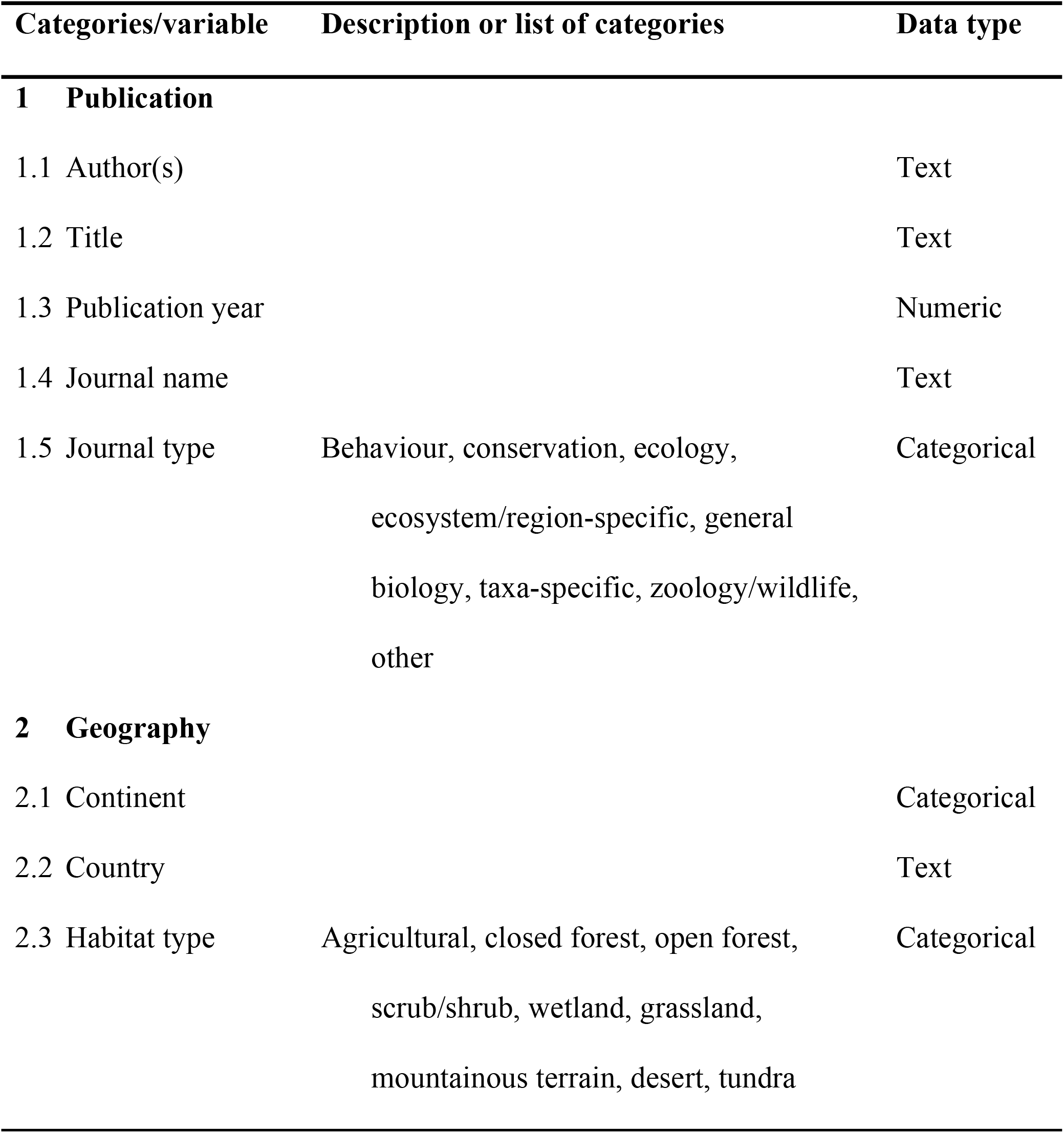
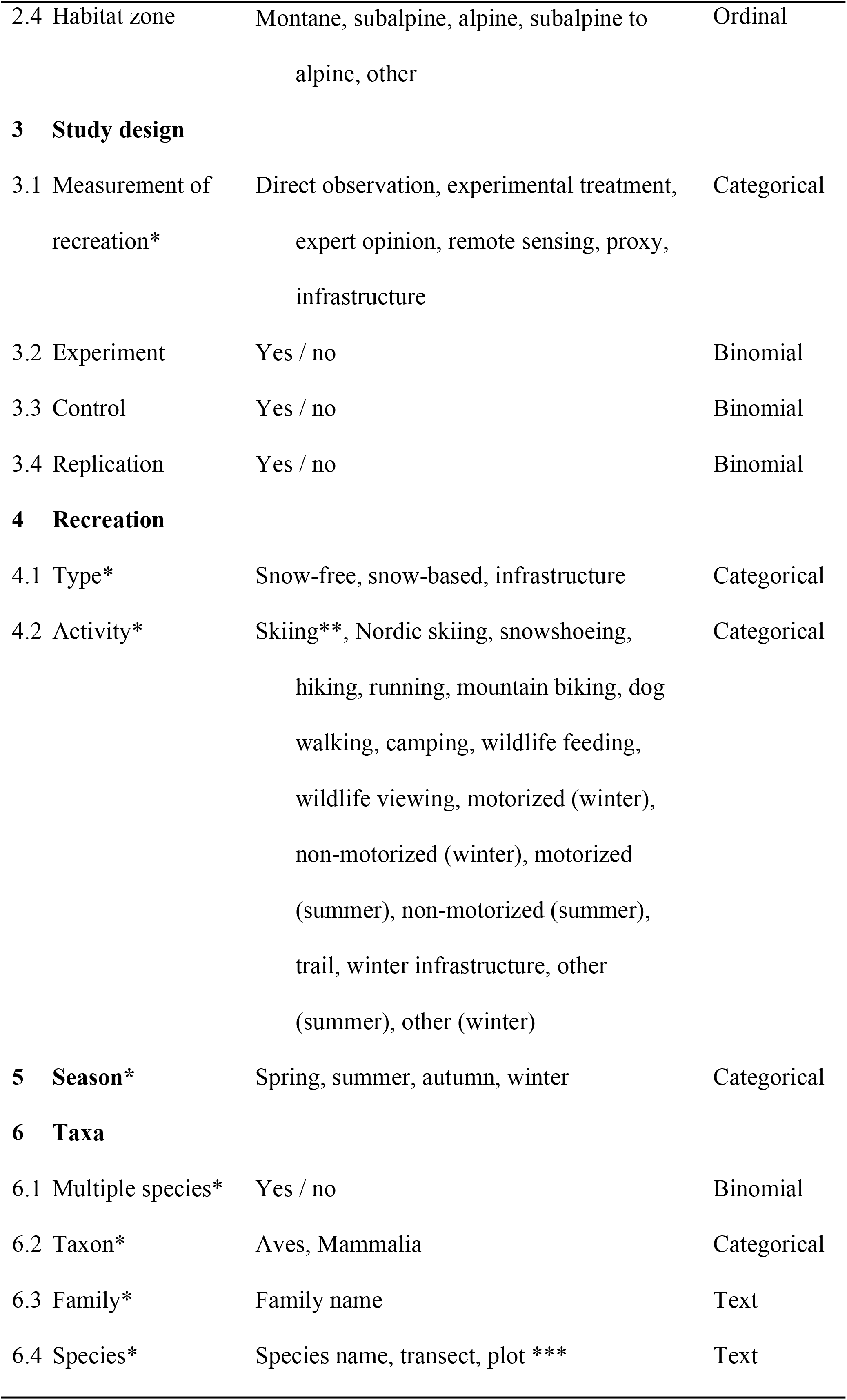
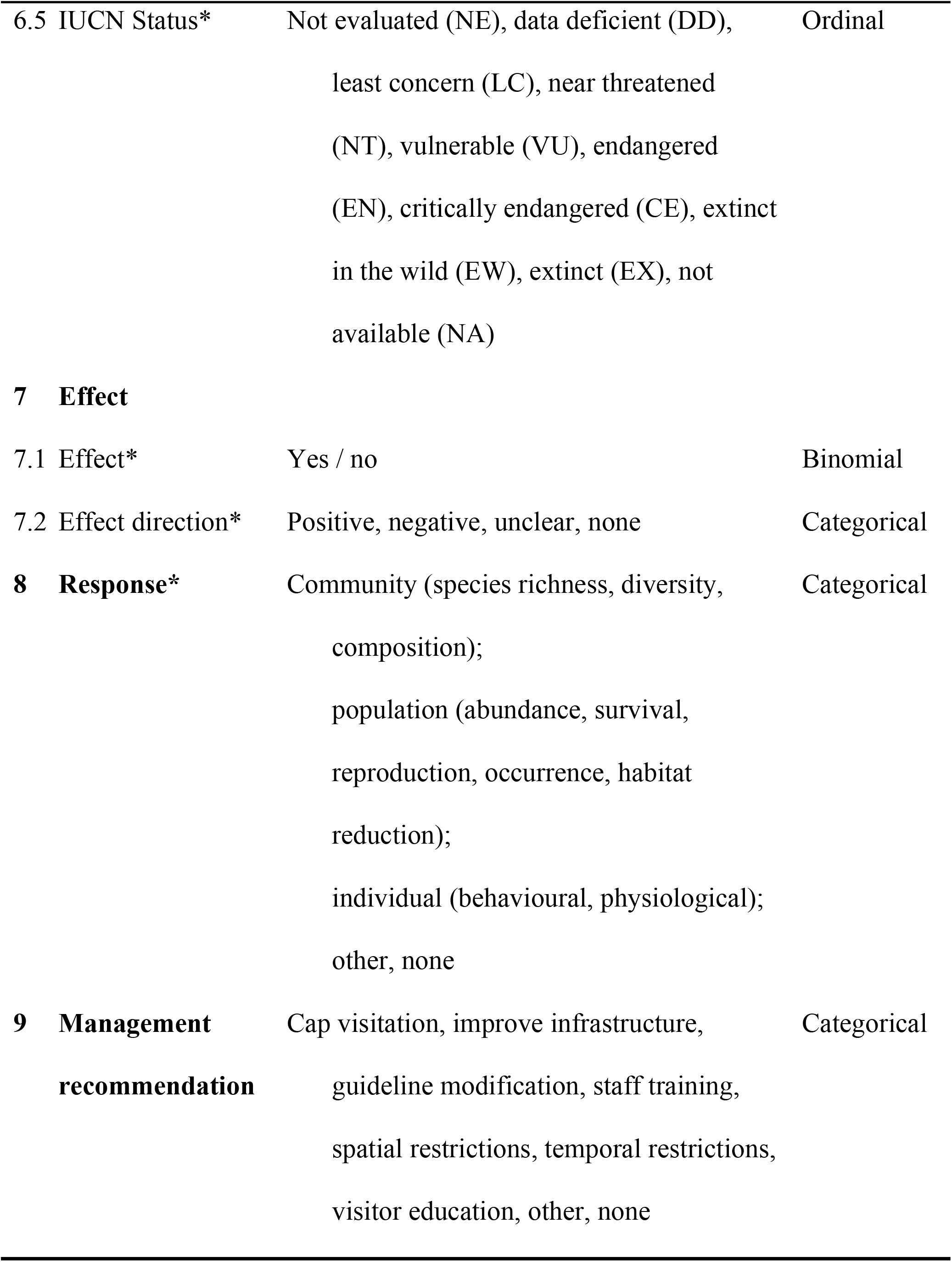
List of catagories/variables collected from the articles included describing the effects of non-consumptive outdoor recreation on mammals and birds in montainous areas. * If an article included more than one example of recreation, seasons, species, effects or responses, each combination was treated as a seperate finding. ** “Skiing” included alpine skiing, backcountry skiing and snowboarding. *** For taxa, we recorded whether more than one species was included in the article or whether a potential multi-species approaches (transect, plot), was used to collect species data. For further details see the S1 Appendix.

**Fig 1:**
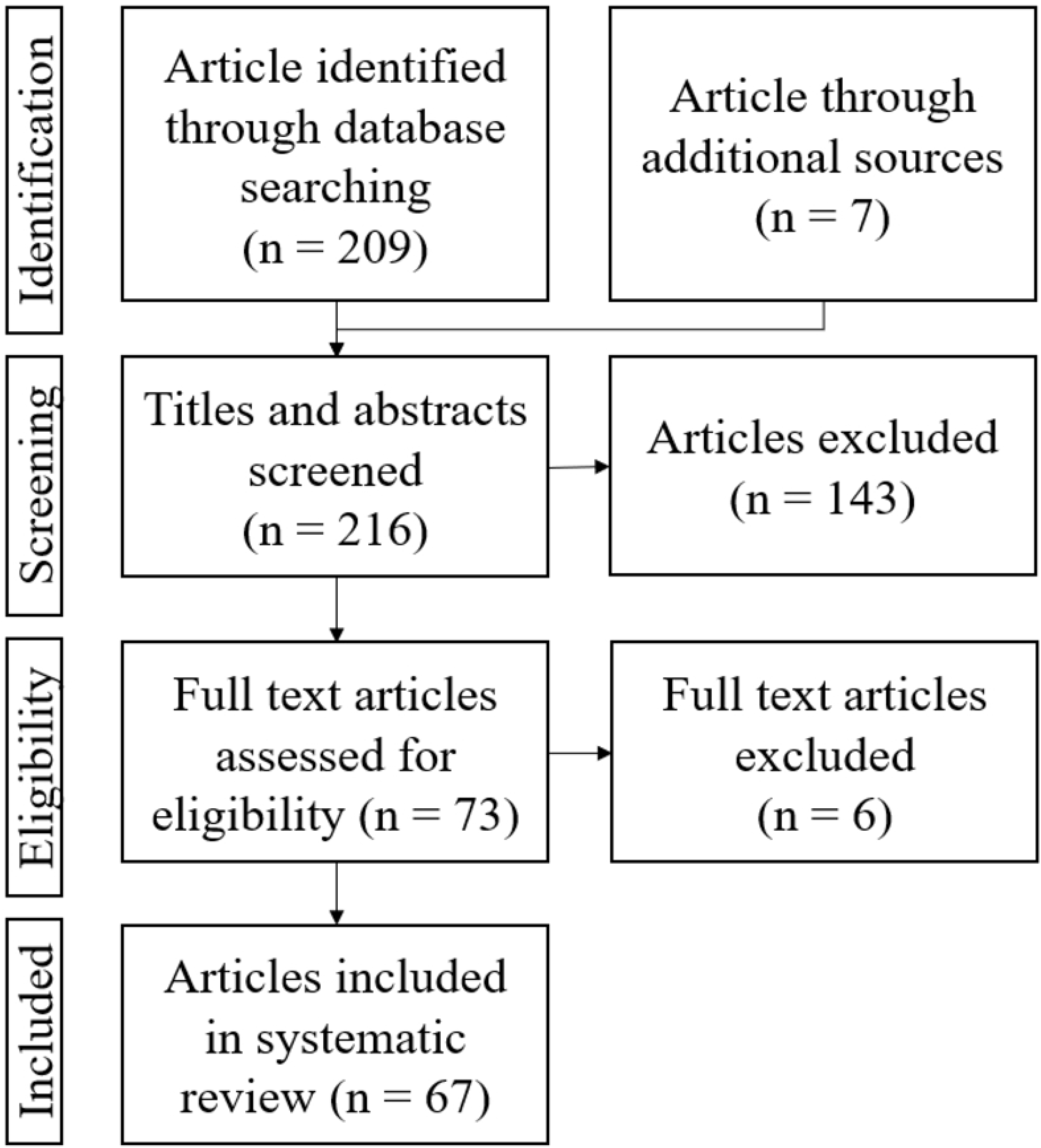
PRISMA literature flowchart. The figure describes the process used for the review including the number of articles identified, retained and discarded in each step adapted from Moher et al. (34). Additional sources are: (15,38–43). For the PRISMA checklist, see S3 Table.

### 2.2 Data collection and processing

Following the PRISMA protocol, we recorded the number of articles in each stage of the screening process and the criteria used to include/exclude them (Fig 1; 52). Initial screening involved reading the titles and abstracts of the articles identified to check which were related to our research questions and criteria. Articles were included if they: 1) addressed the specific topic, 2) focused on mammals and/or birds, 3) discussed an outdoor recreational activity or infrastructure and 4) included research in montane, subalpine and/or alpine habitats. The remaining articles were then read in full to identify those that: 5) included original research on effect(s) of recreation on mammals and/or birds, 6) contained empirical data (e.g. no reviews were included), 7) offered statistical analysis and 8) focused on non-consumptive recreation.

We collected data about the remaining articles in an Excel database (Table) to investigate: 1) who conducted the research, and 2) where the study was conducted, 3) which methods were used, 4) what types of outdoor recreation activities and/or infrastructure were assessed, 5) which season(s) were investigated, 6) which taxonomic classes were studied, 7) which effects were found, 8) how animals reacted, and 9) what management recommendations were made. To facilitate comparison with previous reviews, we used similar categories/variables to those in Larson et al. (30) and Sato et al. (18). Where articles described multiple measurements of recreation, recreation types, seasons, species, effects or responses, these variables were entered in the database as individual results. We treated each study and result as equally valid. Details of the criteria used for the categories are described in the S1 Appendix.

### 2.3 Reporting and statistical analysis

Analysis of the database was conducted using the R Software (Version 1.2.5042; RStudio Team 2020) with data preparation, summarising and visualisation using the “tidyverse” package (45). First the number of publications per year and journal type were quantified. To describe the geographic distribution, we reported the number of articles per elevation zone and continent and visualised the number of articles per country using the package “sf” (46). To examine if the supply of literature related to potential demand, we compared the number of articles per country with the countries ruggedness index, which quantifies the topographic heterogeneity in wildlife habitats (47; we grouped the continuous variable into five equal sized groups for the statistical tests, see below). We also tested if the distribution of the species investigated was proportional to what would be expected based on species diversity per taxonomic family according to the IUCN (2020). We then formally tested whether the distribution of research (supply, observed) differed from expected (demand) using Chi-square Goodness of Fit tests for small sample sizes (49,50) and Fishers exact tests (package “stats”; 51). As Larson et al. (30) stated, the assumption of independence would be violated if an article was assigned to multiple recordings (e.g. several species described in one article). Therefore, we conducted the first test at article level (one article equals one recording) while for the second all articles that accounted for more than one result in the database were excluded.

Next, we summarised articles with specific study design and how recreation was measured. We then analysed which outdoor recreation activities had been studied and how often. To assess the volume of evidence for a specific type of activity affecting mammals or birds, we estimated the overall percentage of results that were statistically significant and showed effect directions (negative, none, positive, unclear). We also did this for season. In a next step, we summarized taxonomic groups (vertebrate class, family and species) to identify which taxa were investigated, how often and where there was evidence of negative effects from recreation. This included assessing the conservation status of species including if they were threatened (48). To evaluate the responses of specific species, we analysed responses by season and recreation activity. Moreover, we visualized response types per recreation activity. Finally, we analysed the proportion of articles that include management recommendations. We considered p-values < 0.05 as statistically significant.

## 3 Results

### 3.1 Publication characteristics and research designs

A total of 67 articles examined some aspect of non-consumptive recreation on mammals and birds in mountainous areas (for included studies see S2 Appendix). Although not large, there is continuing interest in this topic with an average of 4.8 articles per year. Most articles were published in zoology/wildlife (28.4%) or ecology (22.4%) journals but some were published in taxa-specific (14.9%), conservation (11.9%) or ecosystem/region-specific journals (9%). The most common journals were the European Journal of Wildlife Research (11.9%), Journal of Wildlife Management (7.5%) and Biological Conservation, Ecosphere and the Journal of Applied Ecology with 6% of articles in each.

There were clear hotspots and coldspots in research with most articles reporting on research in Europe (52%; Fig 2a) or North America (37%). Specifically, 32.8% of articles were from the United States of America, 14.9% from Italy, 8% from Switzerland, followed by Poland and the United Kingdom (6% each), Australia, Canada, Germany (4.9% each) and Norway (3%). Nearly half of the research was undertaken in subalpine habitats (49.3%; Fig 2b), followed by alpine (23.9%) and montane habitats (19.4%), although for some articles there was no precise differentiation between these climatic zones (6.5%). Most articles report on research in open forest (37.1%) and mountainous terrain (steep, rugged and predominantly rocky; 23.8%), but also research in grasslands (15.2%). In contrast, scrub/shrub, wetland (6.7% each), closed forest (5.7%) and high deserts (4.8%) were rarely studied. The results were not geographically representative in terms of mountainous area per country, with no significant association between ruggedness within a country and the number of articles from that country (Chi-squared Test, p = 0.741; Fisher’s Exact Test, p = 0.907).

**Fig 2:**
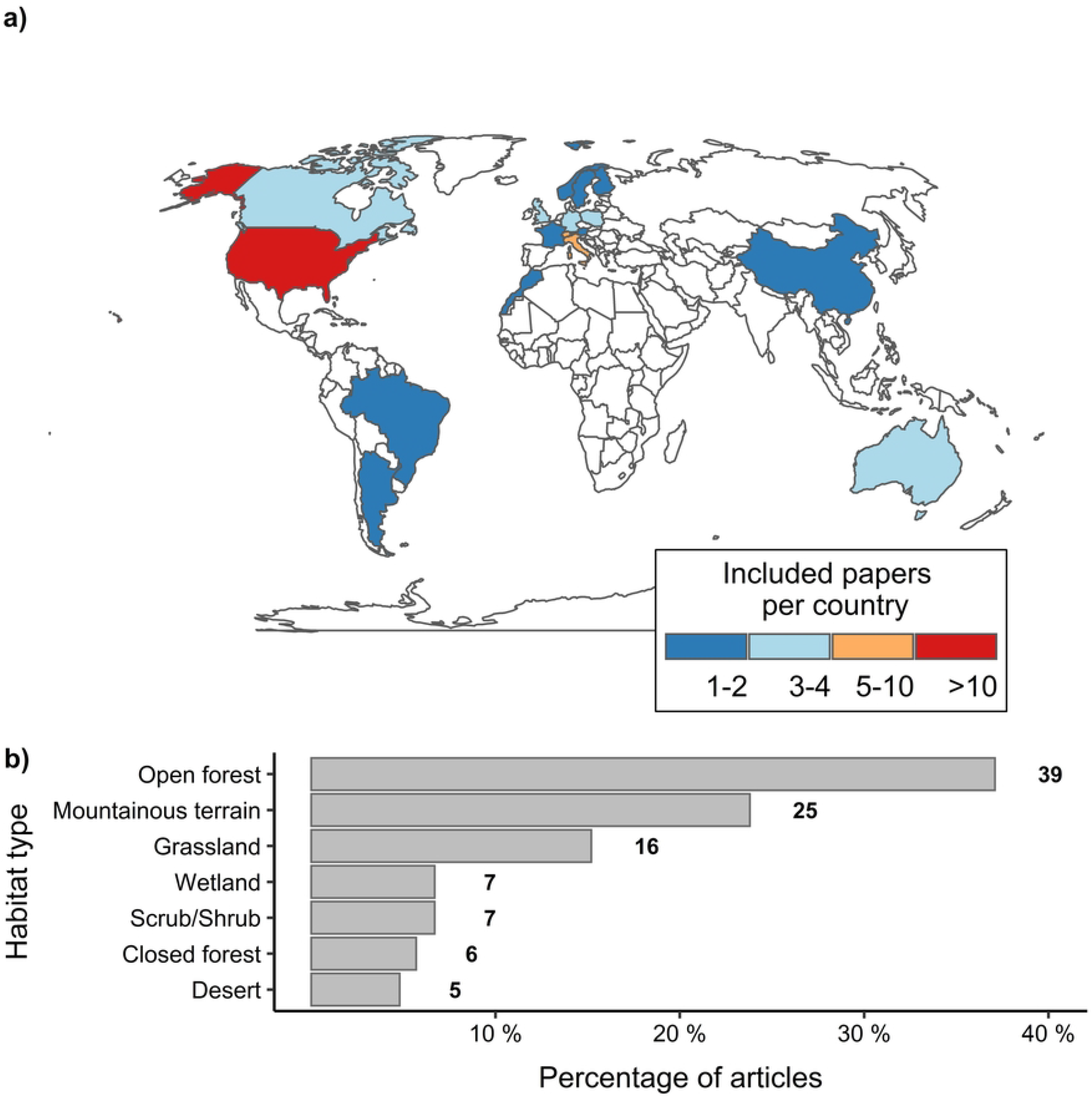
Geographical distribution of articles on the effects of non-consumptive recreation on mammals and birds in mountainous areas: a) including the number of articles per country and b) the distribution of research in relation to different types of habitat, with some articles including research from more than one habitat type. The numbers at the end of the bars give the absolute number of results per habitat type.

In terms of research design, most articles incorporated controls (85.1%) and/or replication (73.1%) of study sites or treatments, but few used experimental treatments (i.e. simulated recreation; 17.9%). Methods used to measure recreation included the presence of infrastructure (52.1%), remote sensing (including automatic counters; 25.4%), experimental treatments (11.3%), proxies for the intensity of usage (5.6%), direct observation (4.2%) or opinions from experts (1.4%).

### 3.2 Impacts of different types of recreation

Most research assessed the impacts of recreation infrastructure for winter activities (23.6%) or trails (19.1%; Fig 4a). There was also research on snow-free (summer) recreation activities (40.2%), mainly hiking or some type of non-motorized activities (i.e. no specific detail of the activity in the article; 11.2%). Where snow-based recreation activities were examined (12.3% of the articles), it was often skiing (6.7%). Reflecting this, most research was done in summer (47.1%) or winter (25.5%; Fig 3) with far less research in spring (15.7%) or autumn (11.8%).

**Fig 3:**
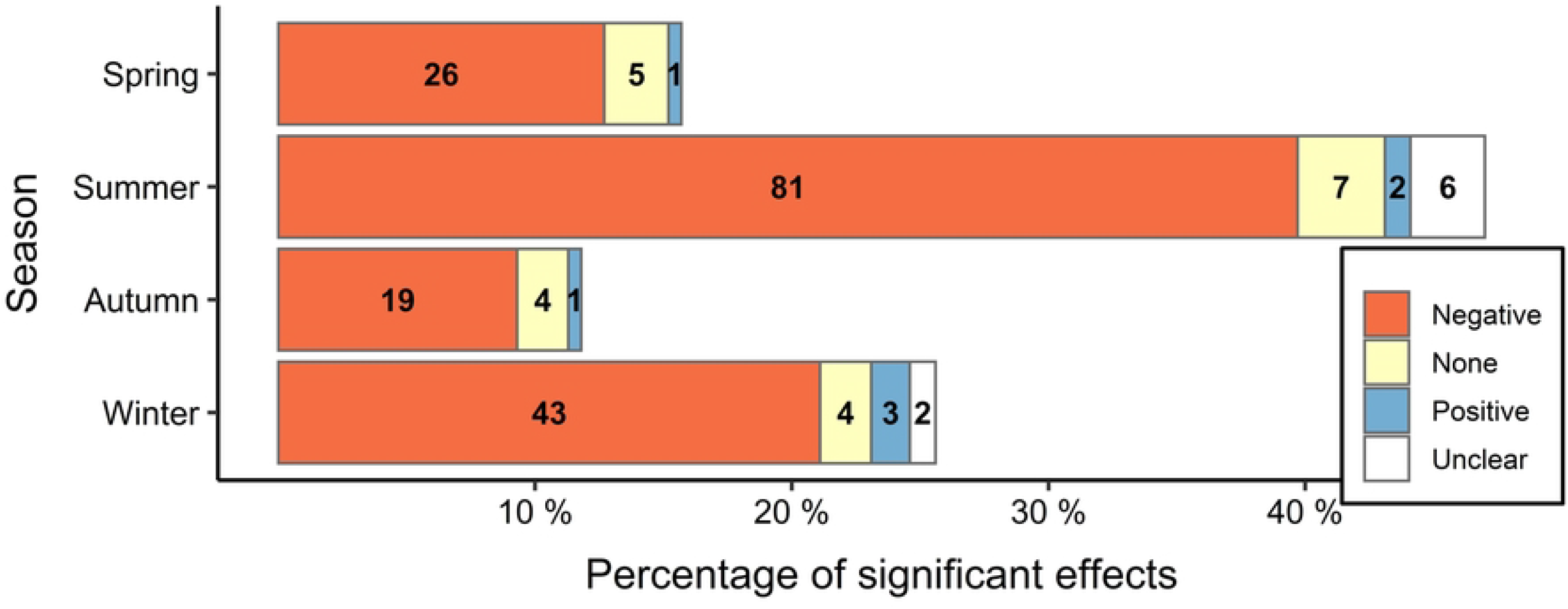
Effects of non-consumptive recreation activities on mammals and birds in mountainous areas by season. Evidence for a negative effect in each season is given by the statistically significant proportion of negative effects compared to all reported effects. The numbers in the bars give the absolute number per category and effect direction.

A total of 90.1% of all articles found some type of environmental effect (Fig 4b), and it was mostly negative (82.2%). This was particularly the case for hiking (negative effects for all 16 articles), skiing (12 out of 13) and non-motorized summer activities (34 out of 37). Only 3.4% of all articles revealed positive effects; with 4 out of 37 articles finding trails had positive effects while just 3 out of 60 articles found positive effects associated with winter infrastructure.

**Fig 4:**
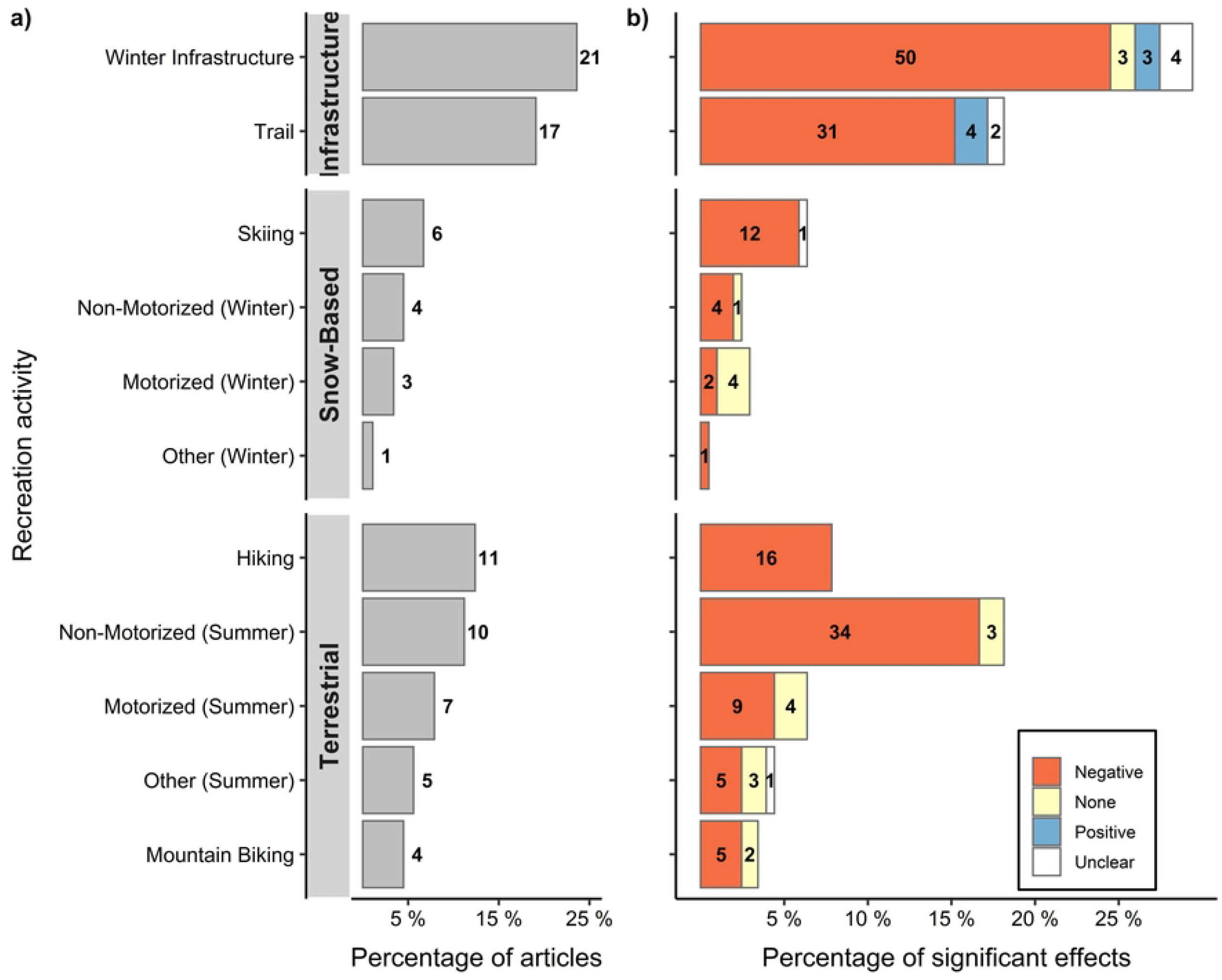
Recreation activities and their effects on mammals and birds in mountainous areas. Including: a) the number of articles that investigated specific activities and/or infrastructure, and b) the percent of results that were statistically significant. Evidence for negative effects of recreation activity were given as the statistically significant proportion of negative effects compared to all reported effects. Some articles accounted for more than one result (e.g. investigated more than one recreation activity and/or infrastructure). The numbers at the end of the bars / in the bars give the absolute number per recreation activity, and effect direction respectively.

### 3.3 Taxa

Research often focused on a single species (77.6%), which was mainly a mammal (69.6%). This research effort is not proportional to species diversity, with no significant relationship between the number of species investigated per family and the number of known species within that family (Chi-squared Test, p = 0.313; Fisher’s Exact Test, p = 1). The species most often researched include red deer (*Cervus elaphus*; 10.8%), black grouse (*Lyrurus tetrix*; 7.5%), bighorn sheep (*Ovis canadensis*), brown bear (*Ursus arctos*) and chamois (*Rupicapra rupicapra*) with 5.3% each and capercaillie/wood grouse (*Tetrao urogallus*; 4.3%). Nearly all of the examined species are on a worldwide perspective considered of least concern (84.3%) according to the IUCN Red List (48), with only 2.9% of the species investigated as nearly threatened, 2.9% vulnerable or 1.4% endangered (Fig 5b). In 8.6% of the results specific species were not mentioned and so their conservation status could not be defined.

**Fig 5:**
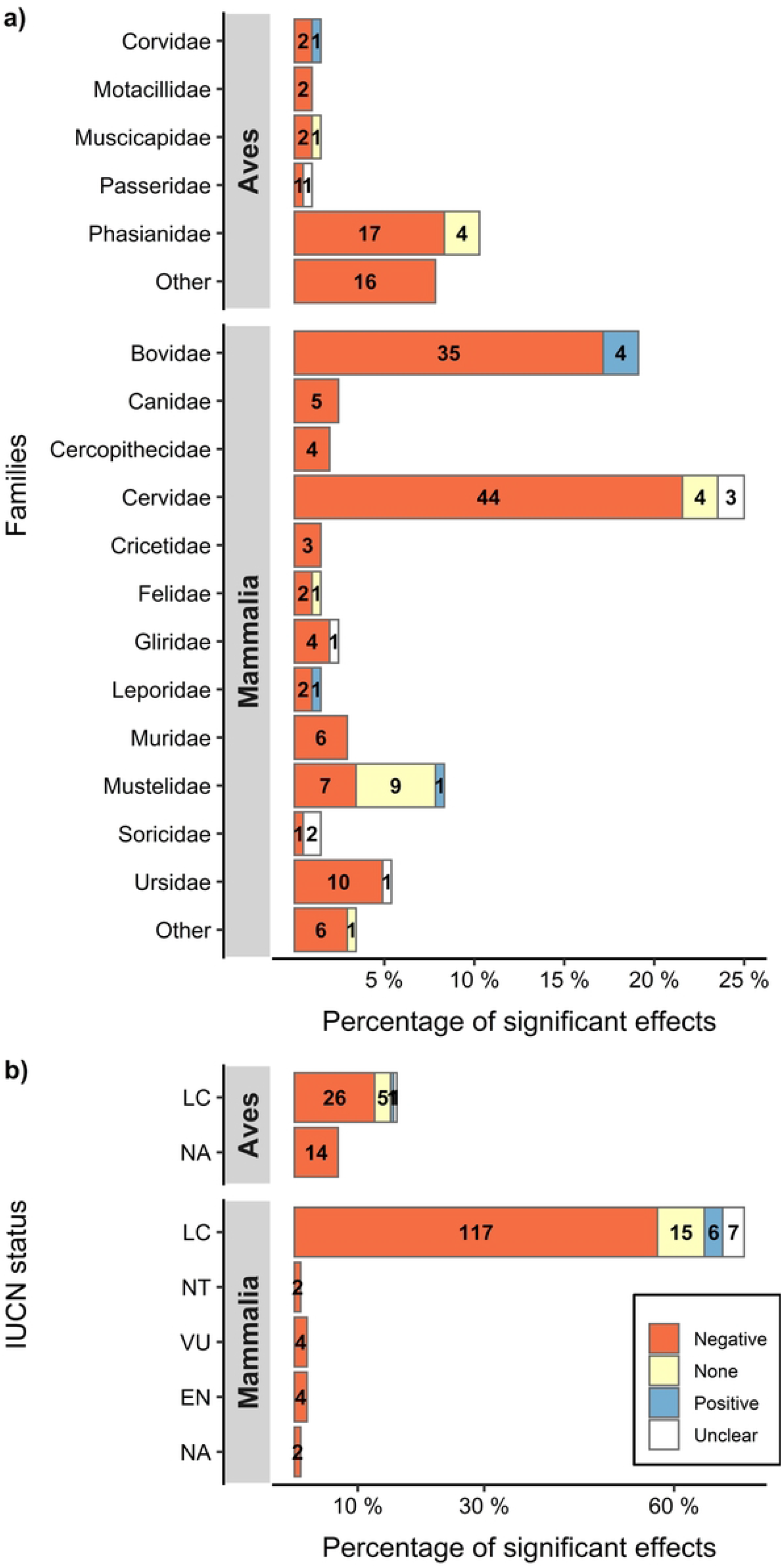
The effects of non-consumptive recreation activities on mammals and birds in mountainous areas by a) taxonomic classes and b) IUCN status. Evidence for a negative effect for each taxon / IUCN status is given by the statistically significant proportion of negative effects compared to all reported effects. The numbers in the bars give the absolute number per category and effect direction. Taxonomic classes with less than two members and/or results of articles that investigated plots or transects rather than species are included in the category “other”.

**Fig 6:**
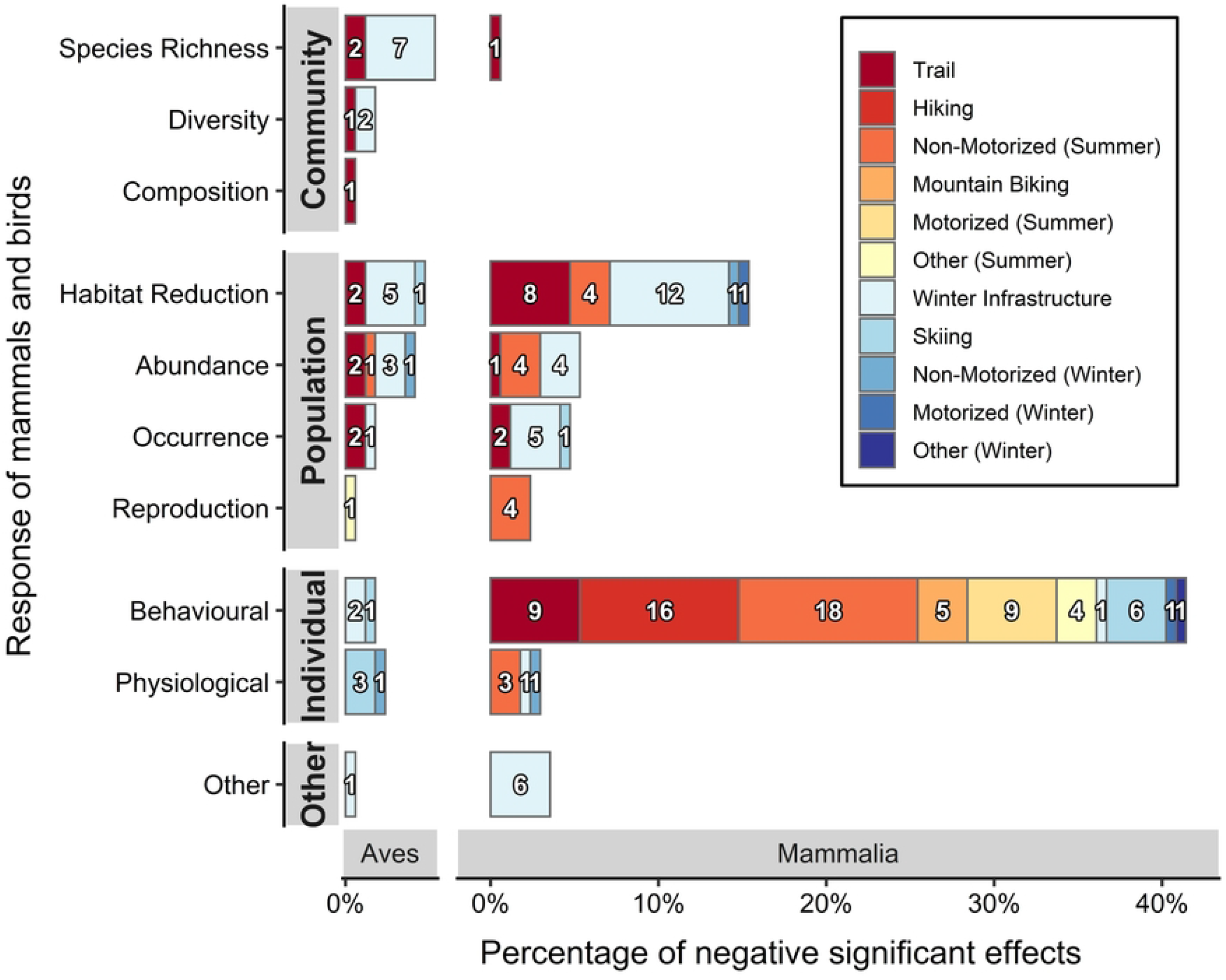
Negative response of mammals and birds in mountainous areas to non-consumptive recreation activities. Red shades show summer, blue shades winter recreation. Some articles reported more than one negative response. From the 204 results, 169 were negative, 20 found no effect, 7 were positive and 8 unclear. The numbers in the bars give the absolute number of negative effects per recreation activity.

Reflecting the focus on relatively few species, most research looked at two families of mammals (cervids, *Cervidae*; 21.6% and bovids, *Bovidae*; 17.2%) and one family of birds (grouse, *Phasianidae*; 8.3%; Fig 5a). Consequently, most negative effects were reported in these families (negative effects: cervids n = 44 of 51, bovids n = 35 of 39, grouse n = 17 of 21). For cervids negative effects were mainly from hiking, motorized activities in summer and mountain biking, with most studies of individuals and documenting changes in behaviour. For example, hikers increased animals vigilance and decreased foraging in cervids (39), while trails used for motorized activities in summer were avoided (64) or resting time decreased when disturbed by mountain biking (62). For bovids, negative effects occurred mostly from trails and non-motorized activities in summer. For example, females in recreational areas had fewer offspring, which led to lower abundance (65). Also group sizes near people were often smaller (66). When positive effects were reported, they were either uncommon or there were few studies in total for that taxa. For example, 4 out of 39 articles on bovids found positive results while for martens (*Mustelidae*) it was one out of 17. For rabbits and hares (*Leporidae*) and corvids (*Corvidae*) 1 out of 3 results reported positive effects. No significant effects of recreation were found for cats (*Felidae*) and flycatcher birds (*Muscicapidae*) in 1 out of 3 results for each. No effects were also found in 9 of 17 results concerning martens, 4 of 21 about grouse and 4 of 51 results concerning cervids.

### 3.4 Response of mammals and birds

Research on community level impacts was uncommon, with very few articles reporting impacts for birds or mammal communities (6.8% of all results; Fig). Where there was research it found reductions in species richness (n = 10 of 10) and/or species diversity (3 of 3), or changes in species composition (1 of 1) due to trails or winter infrastructure. Population level effects (39.7% of all results) were larger than community level effects and were mostly negative as well. This included reductions in habitat (34 of 37), changes in abundance (16 of 21), occurrence (11 of 18) or reproduction (5 of 5). While birds dominated community level studies, mammals did for those assessing population level effects. Trails, winter infrastructure and non-motorized summer and winter recreation were the main causes of negative impacts on populations, with few positive responses reported. Some wildlife had a higher abundance (2 of 21) or occurrence (4 of 18), which was explained as response to trails and winter infrastructure. A total of 42.2% of all results reported effects on individuals and it was nearly always negative. Effects included behavioural (73 of 76), and physiological responses (9 of 10) from a range of activities. Most negative responses were reported for non-motorized winter activities and hiking while few negative responses were documented for trails and winter infrastructure. Finally, 16 results found no response and 7 named other responses of mammals and birds to winter infrastructure.

Changes in individual behaviour had across all seasons the most evidence (i.e. the most number of results), while impacts on abundance and occurrence were only reported in summer (Table 3).

**Table 3:** Response of mammals and birds to non-consumptive recreation in mountainous areas in different seasons, sorted by season and their proportion on all results. Responses with less than 2% are grouped together as “Other”.

| Season | Effect direction | Level response | Response type | N<br>(total = 204) |
| --- | --- | --- | --- | --- |
| Spring | Negative | Individual | Behavioural | 13 |
|  | Negative | Population | Habitat reduction | 6 |
| Summer | Negative | Individual | Behavioural | 34 |
|  | Negative | Population | Habitat reduction | 12 |
|  | Negative | Community | Species richness | 10 |
|  | Negative | Population | Abundance | 9 |
|  | Negative | Population | Occurrence | 5 |
|  | None | None | None | 5 |
| Autumn | Negative | Individual | Behavioural | 11 |
| Winter | Negative | Individual | Behavioural | 15 |
|  | Negative | Population | Habitat reduction | 13 |
|  | Negative | Individual | Physiological | 6 |
| Other | Other | Other | Other | 65 |

### 3.5 Management recommendations

Approximately two third of the articles (72.1%) included recommendations on how to minimize the effects of recreation on mammals and birds (Table 4). Diverse recommendations were most often mentioned (36%) such as habitat conservation and improvement or plans for recreational developments that minimised impacts on populations. While 24% of articles suggested spatial restrictions, there were few articles recommended temporal restrictions (5.8%), caping visitation or visitor education (2.3%). On the other hand, 27.9% of all articles did not contain any management recommendations.

**Table 4:** Summary of suggested management recommendations to minimize the effects of non-consumptive recreation activities on mammals and birds in mountainous areas.

| Management recommendations | Example |
| --- | --- |
| Other<br>(n=31, 36%) | Conservation strategies implemented at landscape scale considering the prevention of new constructions in high conservation value sites (52)<br>Maintenance of vegetation patchiness (e.g. a mosaic of grassy shrubland with scattered trees) along ski runs (53)<br>Enhancing visual protection by maintaining a strip of dense understory along trails (54) |
| Spatial restrictions<br>(n=21, 24.4%) | Prioritize areas for the designation of wildlife refuges with restricted or banned human access (55)<br>Wildlife refuges with a diameter of at least 800 m (54)<br>Reduction of recreation infrastructure density in key habitats (54) |
| Temporal restrictions<br>(n=5, 5.8%) | Strictly regulated nocturnal sport events in areas with sensitive wildlife (15) |
| Cap visitation<br>(n=2, 2.3%) | Limit access by visitors, to small sized groups and low in<br>frequency (56) |
| Visitor education<br>(n=2, 2.3%) | Integrate tourist organizations in the planning process<br>from an early stage and inform recreationists how their<br>activities affect wildlife (54) |

## 4 Discussion

Recreation infrastructure and activities clearly have a range of negative impacts on mammals and birds in mountainous areas. Our review used one academic database, specific search terms and focused on recent research, and hence did not encompass literature only listed in other academic databases, grey literature, research using different terminology and older articles. However, the patterns found often reflected and strengthened those seen in other reviews (18,30), while also identifying new challenges and directions for research. Here we discuss the broader implications from this and previous reviews emphasising key areas for future research.

### 4.1 Where is research focused and where are priority areas for future research?

Although there is continued interest in assessing potential impacts of non-consumptive recreation in mountainous areas on mammals and birds, research remains geographically biased, focusing on Europe (52.2%) and Northern America (37.3%), but rarely Asia, Africa and South America. This is similar to the patterns found by Larson et al. (26; Europe = 26.6%, North America = 37.7%) and Sato et al. (20; Europe = 68.3%, North America = 19.5%). Considering the proportion of alpine and subalpine areas globally, as well as ski areas, South America and Asia remains seriously underrepresented in the literature with limited information about impacts in popular tourism areas in the Andes (29) or the Indomalayan ecozones (31) including China, despite these regions having extensive mountainous areas and their popularity for snow based tourism and recreation (57–59). Biases also occurred in terms of zones and habitats with less research in alpine areas, and in scrub and shrub-land, closed forest and high deserts, agricultural areas or tundra. Although in some situations tourism use of such zones/habitats can be less intense (30), research is still required.

### 4.2 Which activities were studied and where are important gaps?

Currently research remains focused on infrastructure (52.1%), or hiking (12.4%) but not on increasingly popular activities such as skiing (6.7%) or mountain biking (4.5%), mountaineering, climbing and trail running. This is important as without knowledge, management will be less effective and more species and communities harmed. Where there is research on outdoor recreation, negative effects were common. We found that 82.8% of research found negative effects, while Larson et al. (30) found it was 59.4%, Steven et al. (31) found 88% and for Sato et al. (18) 48.8%. In contrast, evidence for positive effects was scarce including for winter infrastructure or trails (here 3.4%), similar to what was found by Larson et al. (26; 14.7%), Sato et al. (20; 17.4%) and Steven et al. (27; 11.6% for birds exclusively).

Motorized activities in summer (30.8%) and winter (66.7%) more often had no effects on mammals and birds than any other type of recreation included. Although the speed of motorized activities can trigger flight initiation distance from further away (26), we and others (26,30) found that non-motorized activities were more disruptive than motorized ones. However, six studies directly comparing impacts on mammals (39,60– 64), found that motorized activities most often triggered similar negative responses to non-motorized ones (behavioural effects or habitat reduction). Considering that motorized activities often use larger areas than non-motorized ones (30) and that direct comparisons between them showed comparable responses, it is probable that the total effects of motorized activities have been underestimated. But again, more comparative research for these and other activities, including the use of proxies and remote sensing techniques to measure recreation, would provide greater insights.

### 4.3 Which mammals and birds were assessed and which need more research?

The focus of current research is not proportional to what would be expected based on species diversity per taxonomic family according to the IUCN (48). This means that the research being conducted predominantly on mammals (69.9%) does not fully match the demand. This accords with Larson et al. (2016; 69.0% of research focusing on mammals) and conservation science in general, whereas research on reptiles and amphibians remains sparse (17). Therefore, research on the effects of recreation on reptiles and amphibians should be a priority as has often been highlighted (30).

Among the mammals most of the research has been on cervids and bovids and found negative impacts providing insights which could be used to better manage activities and minimise impacts for these and other mammals, as shown in chapter 3.5.

Few articles reported positive effects, although in some situations areas near hiking trails had a higher occurrence of bovids (67). This shows how some animals are capable of adjusting to predictable on-trail activities (26,68) with habituation arising more rapidly when stimuli occurred in short intervals and on regularly (69). Trails and winter infrastructure might be such a source of regular disturbance, while hiking and skiing can be off-trail and less predictable (26,68). However, we want to stress that in most cases, infrastructure had negative effects on mammals and birds.

Although grouse are considered to be sensitive to human disturbance (13), and most articles found negative effects from recreation, a few articles found no significant response to recreation. Where no responses were found, it was in response to recreational homes (70), winter infrastructure studied during summer (71), hiking, mountain biking, snowshoeing and backcountry skiing (42). It is worth noting here that some research (42) found that recreation negatively affected the abundance of grouse, but not their occurrence.

### 4.4 Why do we need more applied and management focused research and how can new technology and data sources help?

Based on the current review, it is clear that we need both more research addressing and dealing with management actions, and that research can start harnessing new technology and sources of data to provide important insights including for ecology/social research. There is increasingly broad recognition in environmental management of the need to turn research into ground action, but even for the applied literature reviewed here, not all articles provided actual recommendations for management (72% here and 40% in Larson et al. (30) This is a general challenge where the transition from research to action is often difficult, despite increasing recognition of the need for evidence-based decisions in wildlife and land management, including protected areas (72,73). We therefore urge researchers to specifically address management recommendations in their research using, for example, the approaches Cook et al. (74) presented. We also highlight the need for more empirical research directly evaluating the effectiveness of management actions.

Interdisciplinary approaches combining environmental and social sciences should also be prioritised particularly those evaluating visitor management systems (75). New technology and sources of data are emerging that can be benefit such approaches. For example, researchers are starting to use social science approaches to gain insights into the interactions between backcountry skiers and large mammals (76) including using unmanned aerial vehicles (i.e. UAV, drones; 77,78) or GPS technology to monitor tourists and their impacts. For example, Rupf et al. (16) examined how GPS loggers could be used to determine potential conflict zones involving backcountry skiers and grouse species. Jäger et al. (4) used efficiently crowd-sourced GPS-tracks for a similar analysis. Furthermore, researchers are starting to explore how social media and other sources of user-created data can be harnessed to assess recreational movements (79–82) which could then be used in research into impacts on wildlife.

## 5 Conclusion

Research into the impacts of outdoor recreation on mammals and birds in mountainous areas is available but remains sparse compared to the clear need for data to inform on ground management. With the increasing popularity of outdoor recreation in many mountainous areas, including China and the rest of Asia, as well as South America, more diverse types of recreation activities and changes in recreation during and post COVID-19, understanding how visitors in high conservation value areas affect wildlife and other biota will be even more important. Further research, including the use of new technologies and sources of data, will provide additional insights and facilitate better management of outdoor recreation, the landscapes where they occur and the wildlife they affect.

## Acknowledgements

For the inspiration on how to collect variables we thank Courtney Larson, Chloe Sato and their colleagues.

## Supporting Information

**S1 Appendix**. Details of the criteria used to categorise the content of the original research articles included in the database.

**S2 Appendix**. Articles about recreation effects on mammals and birds in mountainous areas included in the literature review.

**S3 Table**. PRISMA checklist.

## Notes

### Competing Interest Statement

The authors have declared no competing interest.

